# Bacteriophage resistance evolution in a honey bee pathogen

**DOI:** 10.1101/2024.07.09.602782

**Authors:** Emma K. Spencer, Yva Eline, Lauren Saucedo, Kevin Linzan, Keera Paull, Craig R. Miller, Tracey L. Peters, James T. Van Leuven

**Author notes:** Corresponding authors –.

## Abstract

Honey bee (*Apis mellifera*) larvae are susceptible to the bacterial pathogen Paenibacillus larvae, which causes severe damage to bee colonies. Antibiotic treatment requires veterinary supervision in the United States, is not used in many parts of the world, perpetuates problems associated with antibiotic resistance, and can necessitate residual testing in bee products. There is interest in using bacteriophages to treat infected colonies (bacteriophage therapy) and several trials are promising. Nevertheless, the safety of using biological agents in the environment must be scrutinized. In this study we analyzed the ability of *P. larvae* to evolve resistance to several different bacteriophages. We found that bacteriophage resistance is rapidly developed in culture but often results in growth defects. Mutations in the bacteriophage-resistant isolates are concentrated in genes encoding potential surface receptors but are also observed in genes controlling general cellular functions, and in two cases—lysogeny. Testing one of these isolates in bee larvae, we found it to have reduced virulence compared to the parental *P. larvae* strain. We also found that bacteriophages are likely able to counteract resistance evolution. This work suggests that while bacteriophage-resistance may arise, its impact will likely be mitigated by reduced pathogenicity and secondary bacteriophage mutations that overcome resistance.

## Introduction

Managed honey bees (mainly *Apis mellifera*) pollinate around one-third of the world’s pollinator-dependent crops, making their health critical to our food security and integral to food prices. In total, managed honey bees provide $182-577 billion USD/year in global crop pollination services (1, 2). Unfortunately, beekeepers regularly lose around one-third of their colonies every year to a combination of disease and other stressors (3, 4). Efforts to prevent these losses and replace colonies are costly. Among the diseases that contribute to the operational costs of beekeepers is American Foulbrood (AFB). AFB is caused by the gram-positive, spore-forming bacteria *Paenibacillus larvae*. This disease stands out as particularly devastating because of its limited treatment options. In the United States, veterinarian-prescribed antibiotics can be used to clear *P. larvae* infections but the emergence of antibiotic-resistant strains of *P. larvae* raises concerns over AFB management (5–8). Moreover, antibiotics may not eliminate *P. larvae* spores, which can remain in the hive for decades (9–11). In many European Union countries, regulations over antibiotic use and the level of antibiotic residues in bee products curtails their use. As a response to this crisis, researchers have begun to investigate the use of bacteriophages—viruses that specifically target and infect bacteria—as potential allies in the fight against AFB (12–18). These bacterial predators hold immense promise due to their precision, efficacy, and eco-friendly nature (19, 20). Bacteriophages (phages) present an attractive solution to the problem of antibiotic resistance.

Many phages that infect *P. larvae* have been isolated, and attempts have been made to classify them according to the ever changing universal viral taxonomy (https://ictv.global/about/charge) to the genus and species level, such as those phages included in the genuses *Fernvirus, Vegasvirus and Halcyonevirus* (Figure 1) (12, 13, 15–17, 21–23). These phages can kill nearly all known *P. larvae* genotypes in culture (12, 21) and reduce disease burden when tested on bee hives (17, 24). However, all discovered *P. larvae* phages are temperate, a potentially problematic property for their use as therapeutics because of lysogeny (25) and the ability of temperate phages to transfer genetic information between hosts by generalized and specialized transduction (26, 27). The host range (breadth of susceptible host genotypes) of *P. larvae* phages is very broad—a positive characteristic for their utilization in treating infected hives. Thus, phage cocktails comprised of only a few phages can be designed to treat all or nearly all *P. larvae* genotypes that currently infect *A. mellifera*. The genetic diversity of *P. larvae* genotypes is organized by clustering variants into five subgroups based on enterobacterial repetitive intergenic consensus (ERIC) amplification patterns (28, 29). These subgroups differ in their geographic distribution, prevalence, and infection characteristics. ERIC I and II are currently the most widespread and problematic (30–32).

**Figure 1.**
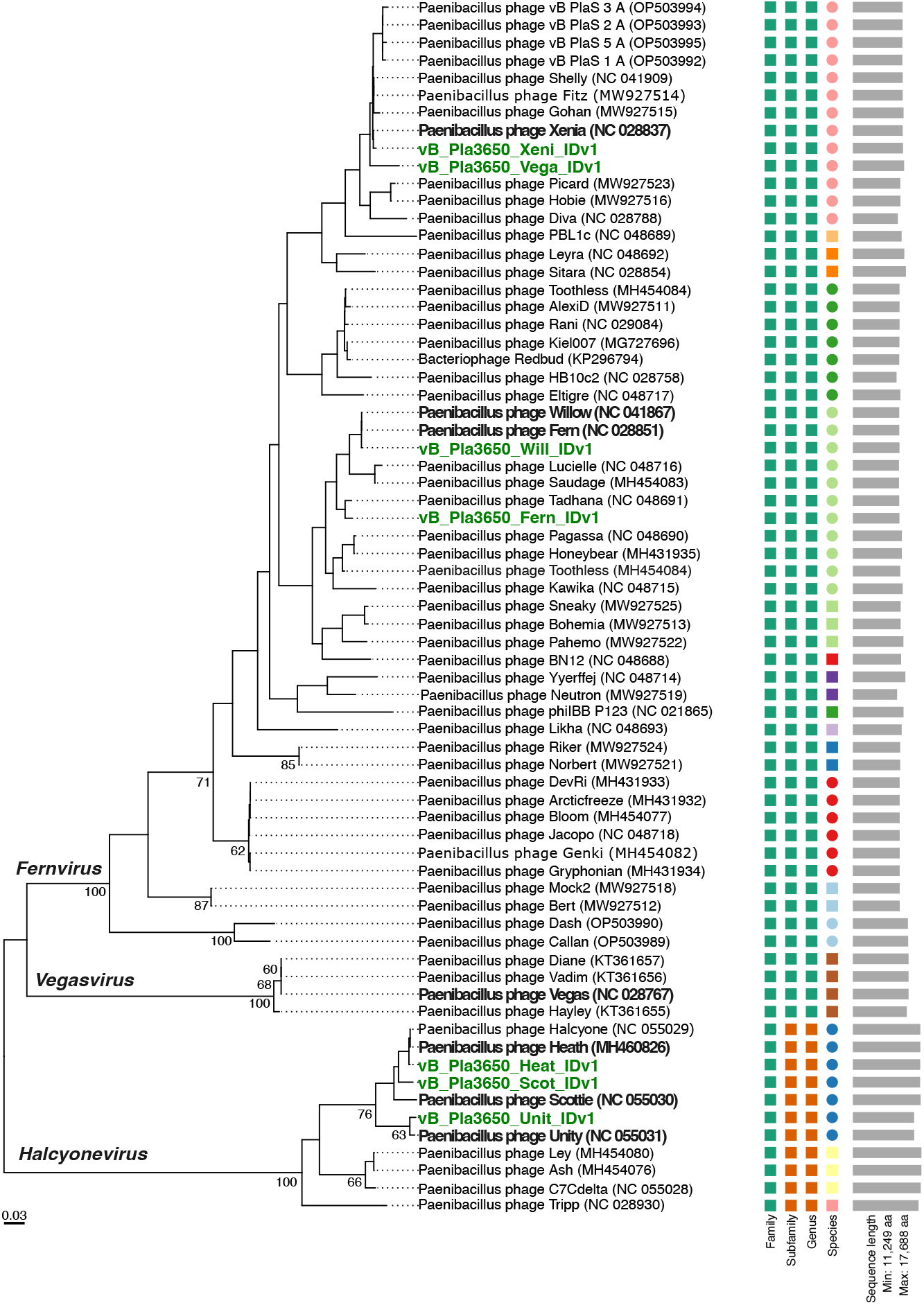
Phylogenomic GBDP tree inferred using the D6 formula based on whole genome amino acid sequence pairwise comparisons, with average support of 31%. The numbers above branches are the pseudo-bootstrap values from 100 replications, using the VICTOR workflow. ICTV genus designations are listend on respective branches (*Fernvirus, Vegasvirus, Halcyonevirus*). Phages used in this study are emphasized with green bold text. Parent phages they are derived from are emphasized in black bold text.

In this study, we assessed the evolutionary capacity of an ERIC I *P. larvae* isolate NRRL B-3650 to gain resistance to phages and identified the genetic determinants of this phage-resistance.

In addition, we assessed how these genetic mutations affected host growth rate, and analyzed how the acquisition of resistance to one phage affects an isolate’s susceptibility to other phages. This study will help us design effective phage cocktails and understand potential problems when using phage therapy to treat infected animals.

## Results

### Re-sequencing of host strain B-3650 and seven *Paenibacillus phages*

We first sought to establish a reliable reference genome for host strain B-3650. To accomplish this, we used both long and short read data to create a hybrid assembly. This approach resulted in an assembly with one complete contig of 4,355,922 bp that was then annotated using Bakta.

All seven phages used in this study were originally obtained from Dr. Penny Amy at the University of Nevada Las Vegas, and were amplified for experimental use on host strain B-3650. In an effort to verify the genetic identity of these phages, we re-sequenced each phage genome used for selection and gave them new naming designations as follows: vB_Pla3650_Vega_IDv1, vB_Pla3650_Fern_IDv1, vB_Pla3650_Xeni_IDv1, vB_Pla3650_Will_IDv1, vB_Pla3650_Heat_IDv1, vB_Pla3650_Scot_IDv1, vB_Pla3650_Unit_IDv1. Phage reads and genomes were submitted to NCBI (see Data availability). We then compared sequence identities between these seven phages and to published Paenibacillus phage reference genomes (Figure 1, Table 1). Whole genome average nucleotide alignments of phages to their respective reference genome showed sequence identities between 99.81% (Heat_IDv1 to Heath) and 89.89% (Fern_IDv1 to Fern), with our phage Vega_IDv1 showing very low identity to phage Vegas, at 29.53%. ANI between phages showed that the phage Vega_IDv1 sample was most closely related to phage Xenia, and Mash distance estimation classified this phage as a *Fernvirus*.

**Table 1.**
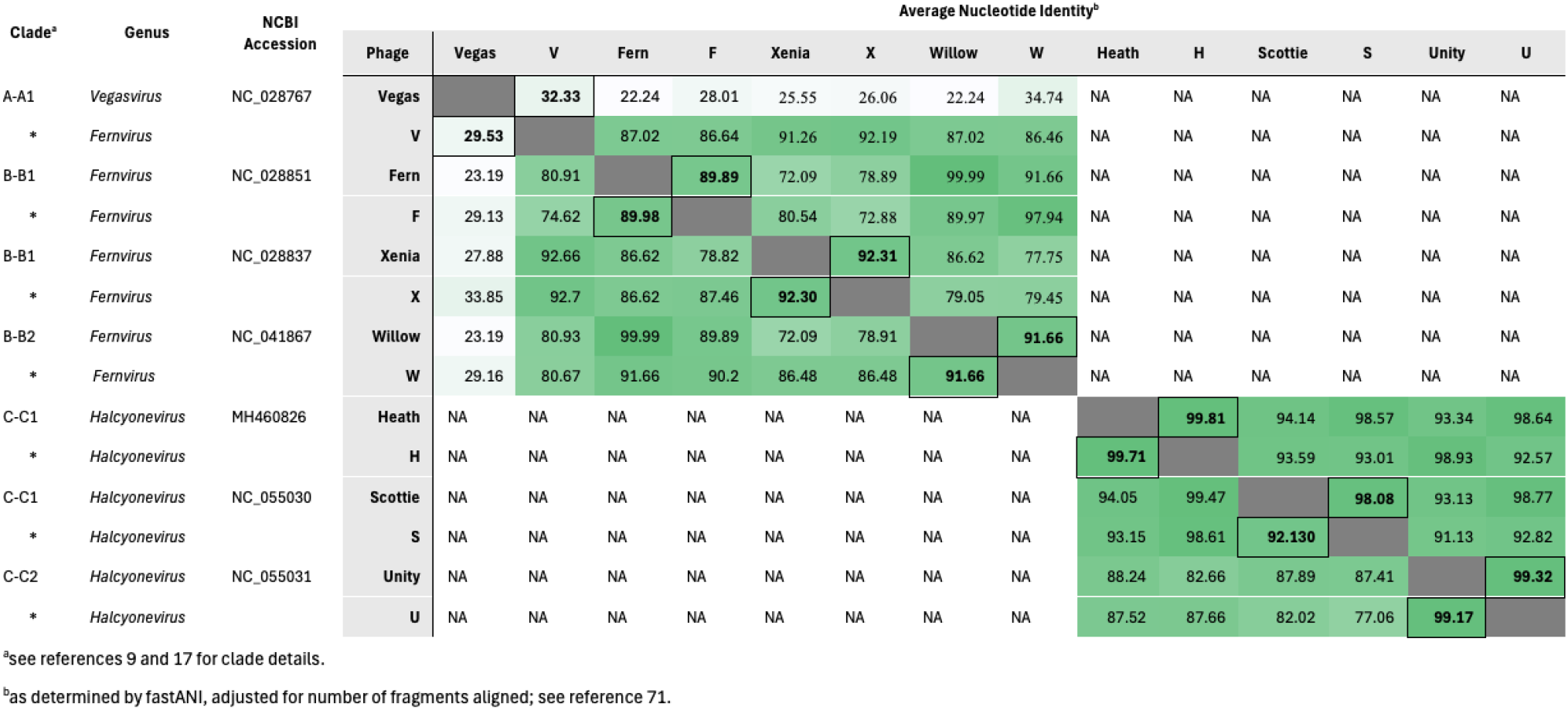
Comparison between *Paenibacillus* phages used in this study (designated by an asterisk and a single letter; e.g. Vega_IDv1 = V) and to previously published reference phage genomes (designated by outlined cells and bold text) using whole genome average nucleotide identity. Lifestyle prediction is shown. Phages were propagated on *P. larvae* sbs*P. larvae* strain B-3650.

### Rapid resistance evolution

To select for phage-resistant isolates, we challenged lawns of *P. larvae* strain B-3650 (ERIC I) with seven different phages (Table 1) at an MOI around 5.

After 24-72 hours, phage-resistant colonies were visible on otherwise cleared plates in all seven challenges. Resistant colonies were rare, with only ~10 colonies (range of 4-114) appearing on plates seeded with 1E6 cells. We picked 3 to 5 colonies from each phage challenge, re-confirmed their resistance to the phage initially used, and sequenced the genomes of 26 isolates. These isolates were named according to the phage that they were challenged with and a one or two letter suffix to denote individual isolates (e.g., Fern_IDv1-resistant isolate yb). In these 26 isolates, a total of 19 unique mutations (13 unique genes) were observed (Figure 2, Table 2).

**Table 2.** Mutations identified in B-3650 phage-resistant isolates selected by seven *Paenibacillus phages*. Superscripts (1-4) indicate different clones (colonies) from one agar plate that initiated the sub-cultures used to evolve resistance.

| Genomic position | Gene name | Predicted Function | Mutation Type | Nucleotide change | Mutation Effect | Number of isolates selected by phage: |  |  |  |  |  |  | Total isolates with mutation: | Isolates IDs: |
| --- | --- | --- | --- | --- | --- | --- | --- | --- | --- | --- | --- | --- | --- | --- |
|  |  |  |  |  |  | Fernvirus |  |  |  | Halcyonevirus |  |  |  |  |
|  |  |  |  |  |  | Vega IDv1 | Xeni IDv1 | Fern IDv1 | Will IDv1 | Scot IDv1 | Heat IDv1 | Unit IDv1 |  |  |
| 303,314 | intergenic |  | SNP | G->A | unknown |  |  |  |  | 2 | 2 | 3 | 7 | Heat_IDv1_x<sup>1</sup>,<br>Heat_IDv1_z<sup>1</sup>,<br>Scot_IDv1_r<sup>2</sup>,<br>Scot_IDv1_t<sup>2</sup>,<br>Unit_IDv1_l<sup>3</sup>,<br>Unit_IDv1_m<sup>3</sup>,<br>Unit_IDv1_o<sup>3</sup> |
| 563,806 | prsA | foldase | ins | +7 bp | FS, early stop |  |  |  |  | 1 |  |  |  | Scot_IDv1_b<sup>4</sup> |
| 563,948 | prsA | foldase | ins | +A | FS, early stop |  |  |  |  | 1 |  |  | 1 | Scot_IDv1_y<sup>4</sup> |
| 564,051 | prsA | foldase | ins | +T | FS, early stop |  |  |  |  |  |  | 1 | 1 | Unit_IDv1_z<sup>1</sup> |
| 564,059 | prsA | foldase | del | -139 bp | FS, early stop |  |  |  |  |  |  | 1 | 1 | Unit_IDv1_ab<sup>4</sup> |
| 564,128 | prsA | foldase | del | -A | FS, early stop |  |  |  | 1 |  | 3 |  | 4 | Heat_IDv1_a<sup>4</sup>,<br>Heat_IDv1_b<sup>4</sup>,<br>Heat_IDv1_y<sup>1</sup>,<br>Will_IDv1_z<sup>1</sup> |
| 1,075,265 | hypothetical | hypothetical | non-syn | A->G | Asn->Asp |  | 1 |  |  |  |  |  | 1 | Xeni_IDv1_z<sup>1</sup> |
| 1,285,883 | gerE | Major transcriptional regulator of sporulation | Ins | phage integration |  | 3 |  |  |  |  |  |  |  | Vega_IDv1_x<sup>1</sup>,<br>Vega_IDv1_y<sup>1</sup>,<br>Vega_IDv1_aa<sup>4</sup> |
| 1,360,187 | lepA | Translation elongation factor | ins | phage integration |  |  |  | 3 |  |  |  |  |  | Fern_IDv1_yb<sup>4</sup>,<br>Fern_IDv1_z<sup>1</sup>,<br>Fern_IDv1_x<sup>1</sup> |
| 1,366,666 | dnaJ | chaperone protein | del | -51 bp | del 17 aa |  | 2 |  | 2 |  |  |  | 4 | Will_IDv1_a<sup>4</sup>,<br>Will_IDv1_x<sup>1</sup>,<br>Xeni_IDv1_Ba<sup>4</sup>,<br>Xeni_IDv1_Bb<sup>4</sup> |
| 1,676,262 | bglG | mannitol operon activator | non-syn | T->A | Phe->Ile |  |  | 1 |  |  |  |  | 1 | Fern_IDv1_yb<sup>4</sup> |
| 1,796,378 | rpiR, mutT, mdtJ, emrE, acrR | tagatose utilization, multi efflux pump, ethidium bromide resistance, transcription | del | -2038 bp | 4-5 gene deletion | 2 |  |  |  |  |  |  | 2 | Vega_IDv1_x<sup>1</sup>,<br>Vega_IDv1_y<sup>1</sup> |
| 2,034,618 | mobile element protein | mobile element protein | non-syn | G->A | Ala->Thr | 1 |  |  | 1 |  | 2 | 2 | 6 | Heat_IDv1_x<sup>1</sup>,<br>Unit_IDv1_o<sup>3</sup>,<br>Vega_IDv1_x<sup>1</sup>,<br>Will_IDv1_x<sup>1</sup> |
| 2,131,639 | cwlJ | spore cortex-lytic enzyme | syn | C->T | NA |  |  |  |  |  |  | 1 | 1 | Unit_IDv1_ab<sup>4</sup> |
| 2,187,569 | gntR | transcriptional regulator | del | -G | FS, early stop |  |  |  |  |  |  | 1 | 1 | Unit_IDv1_ab<sup>4</sup> |
| 2,269,951 | mobile element protein | mobile element protein | del | -C | FS, early stop |  | 3 | 2 |  | 1 | 2 | 3 | 11 | Fern_IDv1_z<sup>1</sup>,<br>Fern_IDv1_x<sup>1</sup>,<br>Heat_IDv1_z<sup>1</sup>,<br>Scot_IDv1_r<sup>2</sup>,<br>Unit_IDv1_m<sup>3</sup>,<br>Unit_IDv1_o<sup>3</sup>,<br>Xeni_IDv1_Ba<sup>4</sup>,<br>Xeni_IDv1_Bb<sup>4</sup> |
| 2,534,497 | ylzA |  | non-syn | C->G | Arg->Pro |  | 1 |  |  |  |  |  | 1 | Xeni_IDv1_z<sup>1</sup> |
| 2,703,591 | sigK | RNA polymerase sporulation specific sigma factor | inversion |  | disruption of coding region |  |  | 1 |  |  |  |  |  | Fern_IDv1_z<sup>1</sup> |
| 3,287,419 | intergenic |  | ins | +A | unknown |  | 1 |  |  |  |  |  |  | Xeni_IDv1_z<sup>1</sup> |

**Figure 2.**
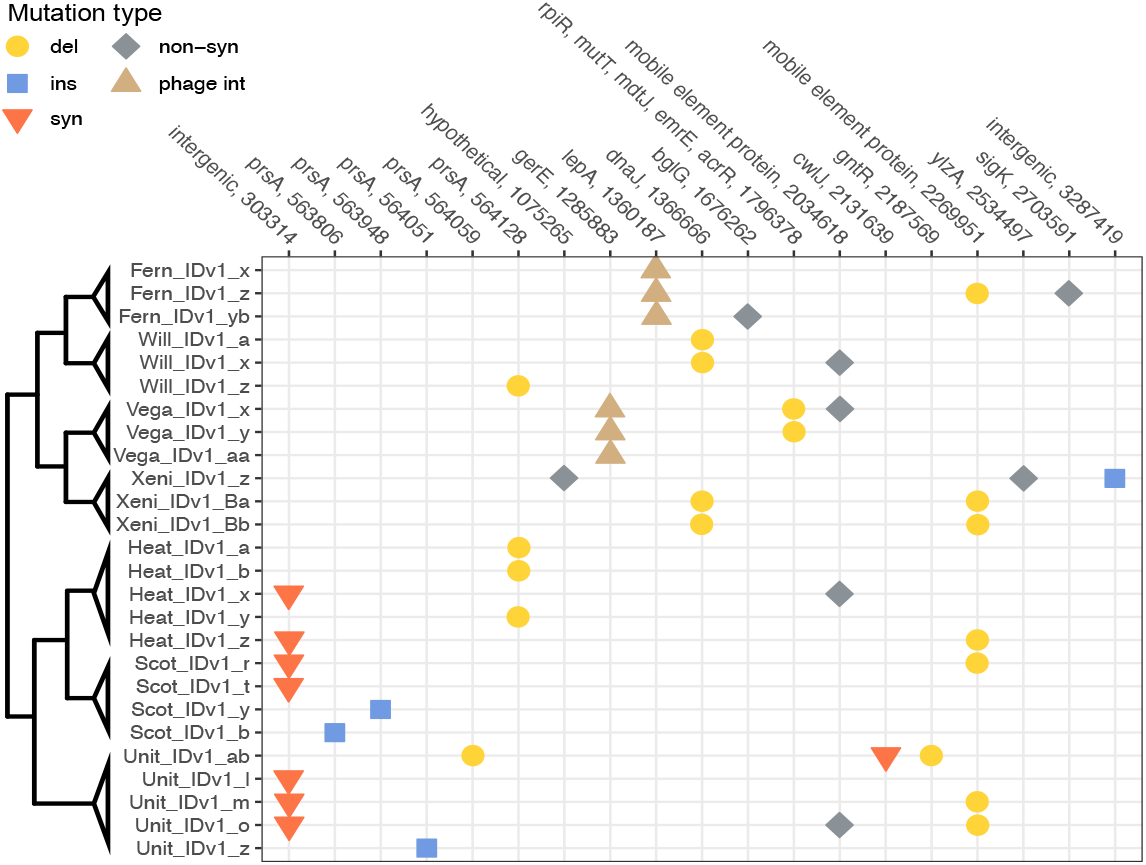
Heatmap depicting the location of mutations in phage-resistant *P. larvae* subp. *larvae* strain B-3650 isolates selected by each phage (Fern_IDv1, Heat_IDv1, Scot_IDv1, Unit_IDv1, Vega_IDv1, Will_IDv1, Xeni_IDv1). Individual isolates are denoted by unique letters following the phage name that they were challenged with. Mutation type is designated as follows: del = deletion (circle), ins = insertion (square), non-syn = non-synonymous mutation (diamond), phage int = formation of lysogen by phage integration (triangle), syn = synonymous mutation (upside-down triangle). Variant analysis was conducted using breseq. The cladogram identifies related groups—branch lengths do not show genetic distance.

Eight samples had mutations in the prsA gene, which encodes for the foldase protein PrsA. For seven of these samples, this was the sole mutation found in the genome. This membrane-bound lipoprotein assists in the folding of secreted proteins (34). The surface exposed region of this protein would be accessible as a phage receptor. Five unique mutations were observed in this gene; two small deletions, one deletion of 138 bp, an insertion of a single thymine, and an insertion of a transposase at base 563,808. This insertion disrupted the coding sequence of prsA. One only isolate with a prsA mutation had other mutations present in its genome (impacting cwlJ and gntR). Neither of these two mutations were observed in any other isolates. Four isolates had a mutation in dnaJ, which encodes for a co-chaperone protein that increases the activity of the heatshock protein DnaK (Hsp70) (35). This protein complex is essential for protein folding and is required for the replication of λ phage in Escherichia coli (36). These four isolates all had the same 51 bp deletion near the N-terminal of the dnaK gene.

This in-frame deletion resulted in the removal of 17 amino acids.

Seven samples had an intergenic mutation (genome position 303,313) in a N-Acetylglucosamine (GlcNAc) biosynthesis gene cluster, just downstream of a predicted transcriptional regulator. GlcNAc is component of the peptidoglycan layer. The genes in this cluster are essential in peptidoglycan synthesis and recycling. Isolate Fern_IDv1_yb had a nonsynonymous mutation in a mannitol operon activator, BglG family CDS. Isolate Xeni_IDv1_z had a nonsynonymous mutation in gene ylzA, which is a regulator of extracellular matrix formation in Bacillus subtilis. This ylzA mutation was accompanied by mutations in a hypothetical protein and an intergenic region. One Unit_IDv1-resistant isolate contained a mutation in cwlJ, a gene encoding a spore cortex-lytic enzyme. This enzyme is involved in peptidoglycan remodeling during spore formation. Two other mutations related to spore formation were observed. One inversion that interrupted sigK, a sporulation-specific sigma factor, and the integration of phage Vega_IDv1 into gerE, a transcriptional regulator of sporulation (this is discussed in a separate section below).

Two isolates that were resistant to phage Vega_IDv1 contained a large deletion of five genes (MurR/RpiR-family regulator, mutT/nudix-family protein, emrE, acrR-family regulator, and a hypothetical protein). MurR/RpiR is a transcriptional regulator of sugar metabolism, including MurNAc synthesis. MutT/nudix-family proteins are hydrolases with broad functions.

*P. larvae* has four genes annotated as mutT/ nudix-family CDS. EmrE is likely a transporter that can provide resistance to ethidium bromide and methyl viologen. The acrR-family gene is likely a transcriptional regulator of efflux pump proteins, possibly involved in biofilm signaling and formation. MurNAc regulation, which may impact peptidoglycan structure, it is unclear how these mutations would provide resistance to bacteriophages.

### Phage Fern_IDv1 and Vega_IDv1 showed integration events in 3650 phage-resistant isolates

Apparent phage integration events were detected in all the Fern_IDv1- and Vega_IDv1-resistant isolates. Evidence for this was first noticed in the breseq (see methods) output as new junctions between the B-3650 and either Fern_IDv1 or Vega_IDv1 phage genomes when breseq was run on a multifasta file containing both genomes. Subsequently, we undertook a careful investigation of potential recombination events by remapping reads and analyzing read coverage. We found elevated (near host depth) coverage for Fern_IDv1 and Vega_IDv1 genomes only in isolates challenged with Fern_IDv1 or Vega_IDv1. Reads mapping to these phage reference genomes matched at 100% identity, suggesting that they were not misaligned. There are regions in B-3650 with high identity to Fern_IDv1 and Vega_IDv1. However, in both cases, large regions of the genome with dissimilarity make it possible to distinguish the phage from the temperate phage regions in B-3650. Using the breseq junction coordinates and read mapping coordinates for soft-clipped reads aligned to the junctions, we identified the integration sites. For Fern_IDv1-challenged isolates, the Fern_IDv1 genome is inserted in B-3650 between bases 1,360,187 and 1,360,222, in the coding sequence for translation elongation factor LepA. These 35 bases are identical to a 35-base region in Fern_IDv1 at genome coordinates 19,500-19,535. This 35 bp region is now duplicated, with copies flanking the now integrated Fern_IDv1 genome. The Fern_IDv1 genome is annotated with a 68 amino acid hypothetical protein in this location, which is directly 5’ to Fern_IDv1’s integrase gene. The Vega_IDv1-resistant isolates now have a phage integrated between genome positions 1,285,883 and 1,285,893. These 9 bases are now duplicated, flanking the now integrated Vega_IDv1 genome, which was disrupted at base 38,418. Integration occurred in the gene for major transcriptional regulator of spore coat formation protein GerE in B-3650 and an intergenic region in Vega_IDv1.

We aligned reads to these new sequences and in both cases, read coverage is even with no breaks, suggesting that these new junctions are correct. No breaks in coverage or soft-clipped reads were found in any isolates other than those challenged with Fern_IDv1 and Vega_IDv1.

### Resistance to one phage changes susceptibility to other phages

To understand how the evolution of resistance will impact treatment with phage cocktails, we characterized the susceptibility of the 26 phage-resistant bacterial isolates to all seven phages used in this study. In all cases, resistance to one phage resulted in cross-resistance to at least one other, typically closely related, phage (Figure 3). This generalization is especially true within the Halcyoneviruses, which are more closely related than the Fernviruses are. For example, isolates resistant to Heat_IDv1, Scot_IDv1, or Unit_IDv1 (Halcyoneviruses) almost always (14/15 isolates) showed some resistance to the other two phages in the cluster. One exception—an isolate that evolved resistance to Scot_IDv1—was completely susceptible to Unit_IDv1. More variation in cross-resistance patterns occurred in hosts that evolved resistance to Fernviruses (Fern_IDv1, Will_IDv1, Xeni_IDv1, Vega_IDv1) (33). Phages Fern_IDv1 and Will_IDv1, which have 99% average nucleotide identity (ANI), usually infected the same host isolates, with two exceptions. Will_IDv1 could infect Xeni_IDv1_isolate_z and Heat_IDv1_isolate_a while Fern_IDv1 could not. These two phage-resistant isolates do not share any mutations. The most distantly related phage in the Fern cluster (Xeni_IDv1, ~75% ANI) could usually (5/6 phage-resistant isolates tested) infect Fern_IDv1 and Will_IDv1 resistant hosts. The opposite was not true, Xeni_IDv1 challenged isolates were usually resistant to Fern_IDv1 and Will_IDv1. Xeni_IDv1 also differed from Fern_IDv1 and Will_IDv1 in its ability to infect other phage-resistant isolates.

**Figure 3.**
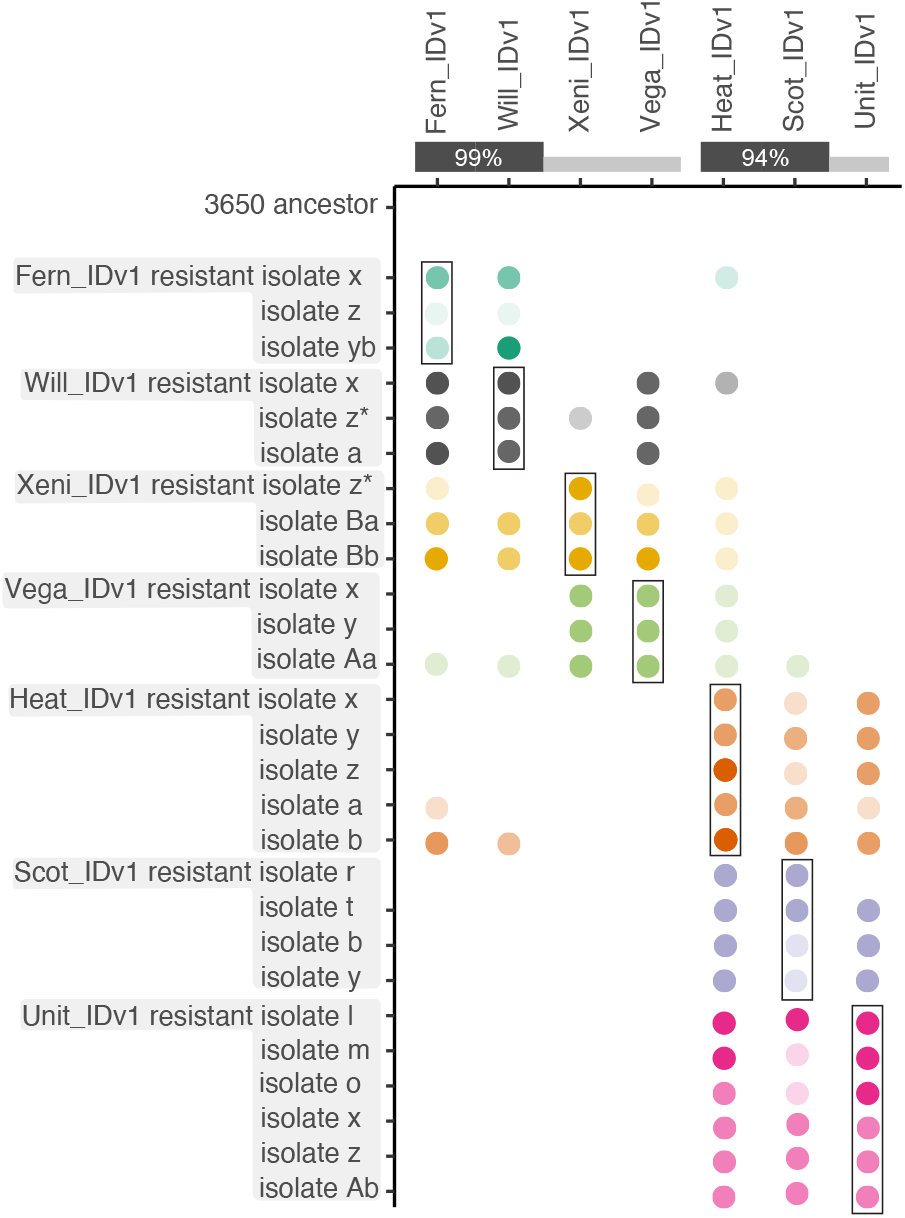
Phage resistance evolution protects hosts against closely-related phages. We tested the susceptibility of all phage-resistant isolates (y-axis) to seven different phages (x-axis). The amount of bacterial grow in the presence of the phages listed along the x-axis is shown by the opacity of colored circles. Dark circles indicate complete or near complete resistance. Partial resistance is indicated by opaque circles. The absence of a circle means that the isolate was killed (no resistance) by the phage named along the top (x-axis) label. The boxes indicate the results of testing a host isolate against the phage that was used to evolve resistance. Nucleotide sequence identity between phages is shown along the top. *Indicates subtle differences in host range among the three methods used. These methods included cross-streaking on agar plates, spot plating phage lysates on bacterial lawns, and soft-agar overlay (see methods for details).

For example, Vega_IDv1-resistant hosts were resistant to Xeni_IDv1, but not Will_IDv1 and Fern_IDv1. While the general pattern is that resistance evolution includes closely related phages, there were many sporadic instances of cross-resistance. For example, Heat_IDv1-resistant isolate b was resistant to closely related phages Scot_IDv1 and Unit_IDv1 as expected, but also to phages Fern_IDv1 and Will_IDv1, which are distantly related. Vega_IDv1_isolate_Aa gained broad resistance to many of the phages. Interestingly, all three Vega_IDv1-resistant isolates were resistant to phages Xeni_IDv1 and Heat_IDv1 which have 92% and <80% nucleotide sequency identity to Vega_IDv1, respectively.

In some instances, resistance was not complete, and occasional plaques were observed on spot and phage overlay plates. To test if resistance was stronger against the challenge phage than against phages that the host acquires cross-resistance to, we compared the efficiency of plating (EOP) of all phages able to grow on a partially resistant host. We found a difference in EOP between these two groups (p=1.5E-3, df=39, paired t-test with Benjamini-Hochberg correction). This result can be visualized in Figure 3 by comparing the transparency of dots along a row to the dot in the box. The transparency of dots in the box are slightly less transparent than dots outside the box. This effect is fairly small, with the average EOP of a phage on a partially resistant host that it was challenged against being 1.6 times lower than other phages that it gained cross-resistance towards.

### Growth defects in resistant isolates

We compared the growth curves of 14 of the phage-resistant isolates of *P. larvae* to the parental B-3650 strain and found that most of the isolates had growth defects when grown without phages (Figure 4, SI Table 4). Lag time, doubling time, and maximum density were calculated and used as response variables in linear models that use either phage or bacterial isolate as predictor variables. All phages except for Xeni_IDv1 caused a significant reduction in maximum density (p<0.05, linear regression). At the level of individual isolates, maximum density differed from B-3650 for 7 of 14 isolates (p<0.05, linear regression). One isolate (Xenia-resistant isolate Bb) had an increase in maximum density compared to B-3650. All other significantly different isolates (Fern_IDv1-resistant isolate yb, Scot_IDv1-resistant isolate b, Unit_IDv1-resistant isolate y, Unit_IDv1-resistant isolate z, Vega_IDv1-resistant isolate x, and Vega_IDv1-resistant isolate y) had reduced maximum growth density. Even within an exposure group—one challenge phage type—there was usually variation in how well different resistant isolates grew.

**Figure 4.**
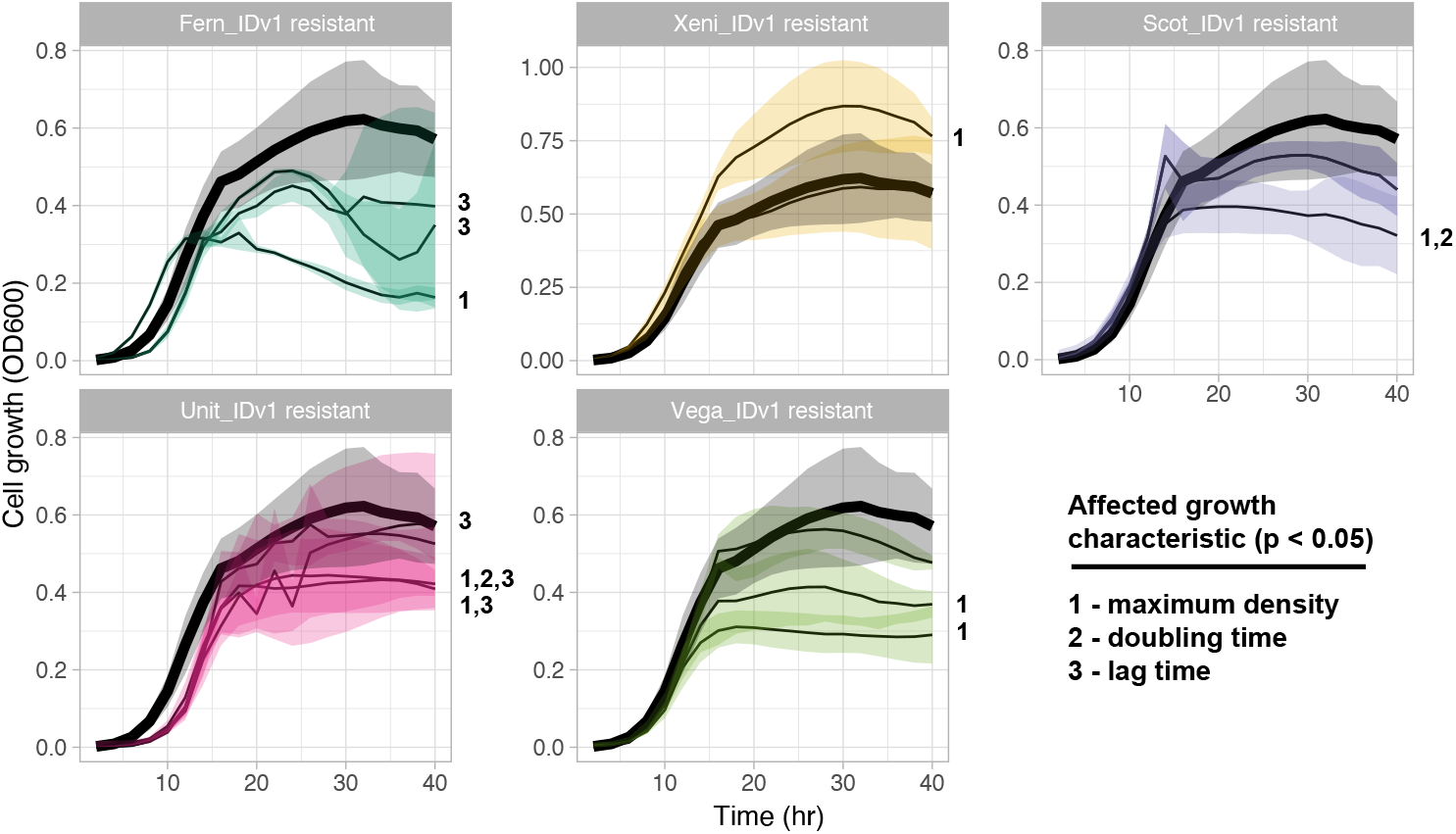
Phage-resistant variants have growth defects. Growth curves of phage-resistant isolates (thin black lines) compared to parental strain B-3650 (thick black line). The standard deviation for at least three replicates is shown by the shaded region. Growth characteristics extracted from growth curves using the gcplyr R package (see methods).

### Potential co-evolution of phages

While verifying the resistance of phage-resistance colonies, we discovered plaques on 18 of the 26 phage-resistant isolates, suggesting that either resistance is partial, or a portion of the phage population can overcome resistance. We further investigated this by measuring the relative efficiency of plating (EOP) of these phages on B-3650 and phage-resistant isolates (SI Table 2). EOP is a commonly used phenotype that is calculated by measuring the number of viruses that can form plaques. In our case, we compared how many plaques form on the ancestor versus resistant hosts. The mean EOP of the initial phage lysates on resistant hosts was 4.5E-5 ± 8.4E-5 (Figure 5). We were unsure if the plaques on resistant hosts were genetic variants with the ability to plaque on resistant hosts or if resistance was incomplete, allowing some phage growth.

**Figure 5.**
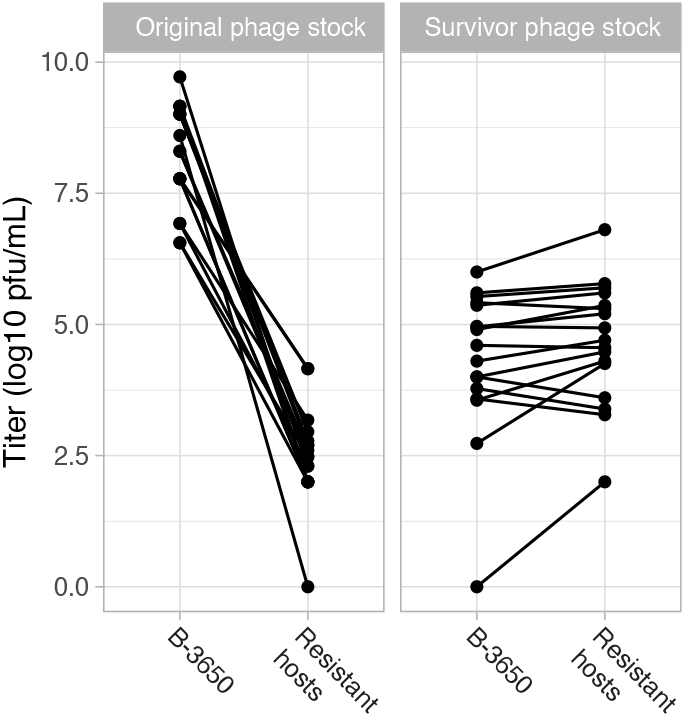
Efficiency of plaquing assays revealed that a small subset of the phages could overcome resistant bacteria. A fraction (18/28) of the phages could form plaques on resistant hosts with very reduced plating efficiency (left panel). When cultivated on resistant hosts, these “survivor” phage isolates had higher plating efficiency on the resistant hosts than the parental host (right panel).

To investigate this, we picked one plaque off every phage-resistant strain and measured the EOP of these picked plaques (titer on resistant host / titer on B-3650). The mean EOP of these plaques was 9.6 ± 24.5 (Figure 5). The minimum EOP for these “survivor” phages was 0.4, meaning that just under half of the phages could plaque on their resistant host after just one round of selection. The EOP of this phage changed from 6.9E-8 before this round of selection to 0.4 after, meaning that only about 1 in every 100,000,000 phages from the original phage population formed plaques on the original host while 4 out 10 could after one round of selection. Therefore, we conclude that rare genetic variants likely exist in the original phage stock that facilitate growth on evolved hosts. These “survivor” phages grew slightly worse (lower titers) on the parental host (Figure 5, p=0.03, df=15, t-test), showing that there is a trade-off associated with growth on phage-resistant hosts.

### Reduced virulence in bee larvae

To determine if the growth defects caused by phage resistance are relevant to bees, we performed a small larvae infection trial with four treatments. Newly hatch larvae were grafted into queen cell cups fed standard feed (see methods) or standard feed with the addition of phage Fern_IDv1 + *P. larvae* strain B-3650, *P. larvae* strain B-3650 alone, or Fern_IDv1-resistant *P. larvae* strain B-3650 (isolate yB). This mutant has one non-synonymous mutation (F109I) in a mannitol operon activator gene and is a lysogen. Because phage Fern was shown to be effective at protecting larvae in a cocktail (see (37)), we only did one replicate (48 larvae) using this phage. The other treatments were replicated at least twice with 12-32 larvae in each replicate. Nearly all (69/72) the mock-infected larvae survived (Figure 6).

**Figure 6.**
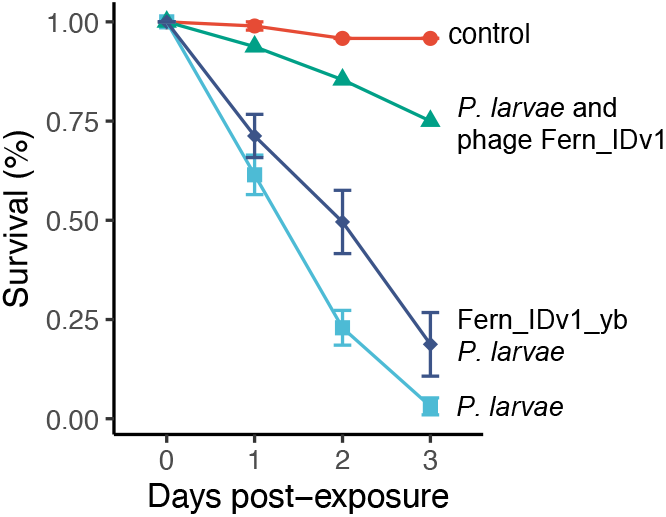
A phage resistant *P. larvae* isolate is less virulent than susceptible *P. larvae*. Recently hatched larvae were reared with diets containing *P. larvae* B-3650, Fern_IDv1-resistant *P. larvae* B-3650, *P. larvae* B-3650 and Fern_IDv1 phage, or media only. 12, 16, or 48 larvae were used in replicate experiments. Standard error bars are show. No replicates were performed for the *P. larvae* and phage Fern_IDv1 treatment (n=48 larvae).

Larvae infected with *P. larvae* and treated with Fern_IDv1 phage survived at ~75%, which is less than control (p=5E-5, Log-Rank test with BH correction, n=1248, df=3). Larvae infected with *P. larvae* strain B-3650 had the worst survival, with only 3% of them surviving 3 days post infection. 18% of the larvae infected with Fern_IDv1-resistant strain B-3650 survived, an 83% increase in survival compared to wildtype B-3650 (p=0.001, Log-Rank test with BH correction, n=1248, df=3).

## Discussion

Using bacteriophages to treat bacterial infections (phage therapy) has remained a promising approach for over a hundred years but much about the basic biology of phage therapy needs to be understood to overcome understandable doubt. Two key questions are addressed in this work; how does phage resistance evolve and how does that resistance alter pathogenicity?

Using the honey bee pathogen *Paenibacillus larvae* and its seven of its associated phages we identified mutations conferring resistance and characterized resulting phenotypic changes in the host.

Phage resistance readily arose in an ERIC-I strain of *P. larvae*, a group of pathogens commonly observed in honey bee colonies (30). Even though *P. larvae* has several anti-phage systems (39), it is susceptible to many previously isolated phages. To ensure that the defense systems in the *P. larvae* isolate we are working with is the same as what was previously characterized, we sequenced the genome of our isolate and ran DefenseFinder (38). We observed no changes in any phage defense related genes and no incorporation of new CRISPR protospacers. When we challenged *P. larvae* with seven different phages, resistant isolates were observable within two days. This type of experimental evolution has been done with many phage-host combinations and is commonly used to identify potential phage receptors (40–44). Each of the 26 resistant isolates that were sequenced had between one and three mutations compared to the ancestor, but only 19 unique mutations were found. Many (10/19) of these mutations were indels, one being a large deletion of five genes. The annotation of many of the affected genes suggest that the mutations have some plausible role in phage receptor presentation; either the genes encode surface proteins or they are involved in sugar or peptidoglycan biosynthesis. It is worth mentioning that all *P. larvae* phage genomes analyzed in a recent study of 48 phages contained a N-acetylmuramoyl-L-alanine amidase (23). These endolysins are likely involved in the lysis of host cells. Thus, is it conceivable that the peptidoglycan mutations we observed somehow alter the cell wall structure, preventing lysis rather than entry.

There was some phylogenetic specificity of resistance mutations. Hosts resistance to phages in the Fernviruses (Fern_IDv1, Will_IDv1, Xeni_IDv1, Vega_IDv1) had mutations impacting dnaJ, a mannitol operon activator, and set of three mutations (hypo, ylzA, intergenic). One isolate (Will_IDv1-resistant isolate z) had a mutation in prsA. The cluster containing Heat_IDv1, Scot_IDv1, and Unit_IDv1 were impacted by five different mutations impacting prsA. Isolates without prsA mutations had a mutation in a transcriptional regulator of N-acetylgalactosamine. It is unclear if the impact of this mutation affects metabolism or cell surface sugar molecule presentation because the genes under control of this regulator region could impact both (45, 46). The conversion of B-3650 into lysogens by Fern_IDv1 and Vega_IDv1 provided resistance against these phages and provides support for the careful use of temperate phages in phage therapy. These phages would be poor choices for treating *P. larvae* infections in honey bee colonies. It is worth noting that B-3650 carries prophages with regions of high sequence identity to Fern_IDv1 and Vega_IDv1, thus it may be advisable to avoid using phages that share sequence homology to prophages in a pathogen’s genome. Four of the six Fern_IDv1 and Vega_IDv1 phage-resistant isolates had additional mutations in their genomes. It is unclear how these mutations contribute (if at all) to resistance.

Phage cross-resistance was largely clustered by phylogenetic relatedness of the challenge phages. Evolving resistance to one phage often confers resistance to phages in the same phylogenetic cluster. There are some interesting exceptions to this rule that are not easily reconciled with the dogma that phages are generally highly host specific. Hosts that evolved resistance to Vega_IDv1 acquired broad protection from phylogenetically diverse phages.

These isolates are all lysogens but isolates x and y have additional mutations including a large deletion of five genes (both isolates) and a non-synonymous mutation in a mobile element protein (isolate x). Others have observed lysogeny providing protection to distantly related phages, however, the mechanism of protection was not resolved (47). Isolates that evolved resistance to Will_IDv1 and Xeni_IDv1 were also commonly resistant to Vega_IDv1. No resistance mutations were found in common between Vega_IDv1 and Xeni_IDv1/Will_IDv1 resistant hosts, so it is unclear how cross-resistance against Vega_IDv1 is gained. Will_IDv1 resistance mutations involve genes dnaJ and prsA.

Several instances of asymmetric resistance acquisition were observed. For example, isolates that gained resistance to Will_IDv1 were not resistant to Xeni_IDv1, but those resistant to Xeni_IDv1 were resistant to Will_IDv1. Isolates resistant to Vega_IDv1 were not resistant to Will_IDv1, but Will_IDv1-resistant isolates were resistant to Vega_IDv1. The Scot_IDv1/ Heat_IDv1/Unit_IDv1-resistant isolates were not resistant to Xeni_IDv1, but Xeni_IDv1-resistant isolates gained partial resistance to these three phages. Others have also reported asymmetric cross-resistance. Among 263 phage resistant isolates of P. aeruginosa, Wright et al. (2018) found isolates to be cross-resistant to 10-80% of the other 27 phages used in the study (48). Cross-resistance is also common when phage receptors are transporters involved in antibiotic resistance (49–51). Gao et al. (2022) reported both asymmetric cross-resistance and somewhat sporadic patterns of cross-resistance in Salmonella enterica (52). We found more effective resistance to the phage that was used during evolution than cross-resistant phages as evidence by lower EOP values for challenge phages than cross-resistant phages (p=1.5E-3, df=39, paired t-test with BH correction). Our results are consistent with results from others showing occasional resistance against phages that are distantly related from the challenge phage (48, 53–54). This suggest that even when combinations of phylogenetically divergent phages are used to treat bacterial infections, there is a chance that universal resistance will evolve. Sequencing the genomes of resistant isolates may provide some predictions on the breadth of cross-resistant. Wright et al. (2018) propose that mutations in potential cell-surface receptors (e.g., LPS) are less likely to confer broad resistance than resistance mutations that change regulator genes (e.g., transcriptional regulators).

In this study we found evidence that phage resistance can come at a cost in terms of growth rate and pathogenicity. However, like other studies have found (55, 56), these costs are not universal—we observed growth effects in culture for half of the isolates that we tested. The one isolate that we tested in bee larvae was worse at killing larvae than B-3650, but more isolates should be tested, particularly those that do not have reduced maximum density in culture.

Many other studies have found that evolution to resist phage infection is accompanied by fitness costs, particularly in the context of a natural microbial community (reviewed in (57)) and thus phage therapy is unlikely to result in widespread resistance against phages (58). Phages that have been preadapted (e.g., “trained”) to overcome bacterial resistance have been shown to reduce the emergence and rise of phage resistant genotypes (59, 60). In our experiments, we observed phages that overcame resistance in 18 of the 26 phage challenges. These phages plaqued on phage-resistant isolates but grew to lower titers on the starting, non-resistant host (Figure 5). Plaquing efficiency assays revealed that a small subset of the phages could overcome resistant bacteria. A fraction (18/28) of the phages could form plaques on resistant hosts with very reduced plating efficiency (Figure 5, left panel). When cultivated on resistant hosts, these “survivor” phage isolates had higher plating efficiency on the resistant hosts than the parental host (right panel). We interpret this reduction in titer as a trade-off for being able to utilize the phage resistant hosts. The number of survivor phages observed is consistent with a hypothesis that these phages overcome resistance via mutation, although there are other potential explanations. Given a dsDNA viral mutation rate of ~1×10-7 mutations per nucleotide per infection cycle (61) and roughly 35-45 kb genome sizes and the 107 phages plated, we calculate roughly 40,000 mutants per plate.

This number is much larger than the number of plaques we observed on lawns of phage resistant isolates. Co-evolutionary arms races between phages and their hosts are common in many systems (57, 62, 63) and thus we would expect the same for *P. larvae* phages. Our results suggest that perhaps a cocktail of preadapted and non-preadapted phages from different phage lineages may best work to treat AFB infections.

## Methods

Bacterial and phage strains and culture conditions. *P. larvae* strain NRRL B-3650 and all seven phages were obtained from Dr. Penny Amy (University of Nevada Las Vegas). Glycerol stocks of B-3650 was streaked on Brain Heart Infusion (BHI) agar plates for the isolation of a single genotype (2X re-streaks). A single colony was used to innoculate 3 mL of BHI liquid media, which was subsequently shaken at 200 rpm at 37° C in 5% CO2 for ~30 hrs to an OD600 of ~0.7. Glycerol stocks of the seven phages were revived by mixing freezer stock with 100 μL of OD600 ~0.7 cells (about 1E7 cfu/mL) in warm mBHI top agar (BHI, 1mM CaCl2, 1 mM MgCl2, 0.7% agar). The warm top agar containing phage and bacteria are gently poured onto a solid agar plate and allowed to try for ~30 min at room temperature at which point they are moved to a 37° C, 5% CO2 incubator for overnight growth. A single plaque was picked and amplified to high titer overnight in 3 mL of mBHI with B-3650. New phage stocks were generated by pelleting cellular debris from spent cultures (1000 x g) then chloroform treating the supernatant (50 μL of chloroform added to 500 μL supernatant). Residual chloroform was removed by centrifuging for 4 minutes at 13000 x g and pipetting off the supernatant containing phages. These stocks were titered and stored at 4° C for immediate use.

Isolation and characterization of phage-resistant mutants. For the resistance evolution experiment, we streaked out a loop-full of glycerol stock B-3650 on BHI agar plates. A colony was chosen for each replicate and grown in 3 mL BHI to an OD600 of ~0.7. Three random marks were placed on the bottom of the agar plate, then 100 μL of cells were briefly vortexed with phage stocks (titers provided in SI Table 1) at an MOI of 5 in mBHI top agar (0.7% agar) and gently poured on solid BHI agar plates. Plates were incubated for 72 h at 37 ° C in 5% CO2. Cleared plates with phage-resistant colonies were visible between 24-72 h and the three colonies closest to the marks were picked. This was repeated at least twice (two starting colonies) for each challenge phage except for Unit_IDv1, which required a third replicate to acquire sufficient plaques. In total, four colonies were used to spread the evolution experiments across more days. Table 1 shows which of the four parental colonies that each phage-resistant isolate originated from. After 72 hours the following colony counts were observed; 4 colonies for Heat_IDv1, 7 colonies for Scot_IDv1, 9 colonies for Unit_IDv1, 114 colonies for Xeni_IDv1. To confirm resistance, individual colonies were picked, grown overnight in mBHI broth and streaked across phage stock (phage streak assay) that was dibbled perpendicular to the colony streak (for example see SI Figure 1). Volumes of 20-50 μL of phage stock was dribbled across the agar plate and allowed to soak into the agar for ~2 minutes. A 10 μL loop-full of overnight culture from putatively resistant hosts was streaked perpendicular to the phage. The B-3650 ancestor was always included as a control. This same phage streak assay was used to determine cross resistance shown in Figure 3. Two additional tests were used for cross-resistance testing. First, we spot plated 5 μL of phage stock on lawns of B-3650 and its derivative phage-resistant isolates. Phage stock titers for the spot plating are shown in SI Table 1. Lastly, phage stocks were titered on every host using the top-agar method described above. The titers were then compared to B-3650 to calculate efficiency of plating (EOP). EOP values were compared using one-sided, paired t-tests in R. For the cross-resistance comparisons, the Benjamini-Hochberg multiple testing correction was applied because multiple comparisons to the ancestor were performed. The full matrix of titers (including survivor phage isolates) on every host isolate are provided in SI Table 3.

Growth curves of phage-resistant isolates. The optical density (OD600) of cultures of 14 phage-resistant isolates of *P. larvae* (six that are resistant to Halcyoneviruses and eight resistant to Fernviruses) were measured in 24-well shaking plate assays in an incubating plate reader (BioTek Instrument, Inc. USA). Control wells included BHI broth control, bacteria control, and phage control. The plates were incubated while shaking for 48 hours with the reading of optical density (600 nm) recorded from each well at the interval of every 15 minutes (SI Table 4). OD600 readings were analyzed in R using the gcplyr (version 1.6.0) and lme4 (version 1.1-33) packages (64). Using the gcplyr package, we extracted doubling time, maximum density, and lag time from the growth data. We compared linear models for these three metrics that included the following predictor variables: phage type, isolate, and mutated gene using Akaike information criterion (analysis code is available at https://github.com/jtvanleuven/plarvae_resistance). This analysis identified all three predictor variables as having a significant impact on at least one of the dependent variables. Thus, we performed three ad-hoc linear models to identify which isolates significantly different from B-3650.

Sequencing and analysis. Overnight cultures of strain B-3650 and phage-resistant isolates were harvested by centrifugation and resuspension in lysis buffer followed by DNA isolation (ZymoBIOMICS). Phage whole-genomic DNA was extracted using the Norgen Phage DNA Extraction Kit (optional protease and second elution steps were included). Phage and host strain B-3650 gDNA were submitted for sequencing on an Illumina HiSeq platform (150PE) with Omega (Norcross, GA). B-3650 gDNA was also prepared for Nanopore sequencing using Native Barcoding Kit 24 V14 (SQK-NBD114.24) and sequenced on a MinION device. Bases were called using dorado (dna_r10.4.1_e8.2_400bps_sup@v4.2.0, https://github.com/nanoporetech/dorado) and demultiplexed using guppy (https://nanoporetech.com/document/Guppy-protocol). Illumina sequencing of bacterial strain B-3650 as a control, and 26 phage-resistant isolates was performed by SeqCoast Genomics (Portsmouth, NH). Samples were prepared for whole genome sequencing using an Illumina DNA Prep tagmentation kit and unique dual indexes and sequenced on an Illumina NextSeq2000 platform using a 300 cycle flow cell kit (150PE).

Illumina reads were quality filtered and trimmed using fastp (v0.23.4). Nanopore reads were filtered using filtlong (v0.2.1) (https://github.com/rrwick/Filtlong). Unicycler (v0.5.0) was used to assemble phage genomes using the short read approach with default parameters. Where necessary, phage samples were subsampled to achieve a depth of 100x prior to assembly using seqtk version (1.4-r122) to achieve complete assemblies. A hybrid assembly was produced using Unicycler (65) for strain B-3650 and was annotated using Bakta (66).

This assembly served as the reference genome for variant analysis. Our hybrid assembly of strain B-3650 was compared to strain NZ_CP019651 by pairwise alignment. One single nucleotide polymorphism at genome position 365,471 and one single base indel at genome position 2,269,949 compared to CP019651 were identified. These mutations are in a putative transposase and mobile element protein, respectively. Variant analysis of phage-resistant isolates was conducted by mapping Illumina reads to the hybrid B-3650 reference genome using BREseq v0.38.1 and visualized using gggenomes (67). Custom R scripts (https://github.com/jtvanleuven/plarvae_resistance) were used for data processing and plotting. Additional mapping was performed using bwa mem (0.7.18-r1243-dirty) and visualized in Tablet (v1.21.02.08).

Phage genomes were annotated using pharokka (68) and assembly statistics were generated using bbmap and samtools. Whole-genome based phylogeny of our seven phages and Paenibacillus phages of the genus Fernvirus, Halcyonevirus, and Vegasvirus was conducted using the VICTOR pipeline (69). Previously published phage genomes were first downloaded from NCBI and re-annotated using pharokka, then submitted to VICTOR whole genome amino acid analysis. Whole genome average nucleotide identity calculations were made using fastANI (70) which reports average nucleotide identity and the number of aligned fragments. We adjusted the ANI score by multiplying by the fraction of genome aligned.

Virulence assay. Age-matched 1-3 day old larvae were transported from our apiary to our laboratory in a pre-heated foam nuc box containing a bottle of hot water and were covered with a damp paper towel. Within 30-60 minutes from being removed from the hive, larvae were grafted using a Chinese grafting tool into brown queen cups in 48 well tissue culture plates containing 20 μL of preheated artificial food “A” (44.25% royal jelly, 5.3% glucose, 5.3% fructose, 0.9% yeast extract, 44.25% water).

Larvae were incubated for 48 hours (see conditions below), then 20 μL of artificial food “B” (42.95% royal jelly, 6.4% glucose, 6.4% fructose, 1.3% yeast extract, 42.95% water) was added to cells. Plates were photographed daily and death was measured by observing larval discoloration and the absence of diet consumption. Larvae were reared following standard procedures (71). Temperature and humidity were monitored constantly and remained at 35° C and 90-100% humidity. A dilution series of 10, 50, 100, or 1000 colony forming units in 1 μL of mBHI growth media was added to the artificial food on day zero. Mock-infected larvae had μL of plain mBHI added to their food. Larval survival was compared in R using the “survival” and “survminer” packages. The Cox proportional hazards regression model was fit according to the following formula, (days post infection, survival) ~ treatment. P values were calculated for all pairwise comparisons using Log-Rank tests with Benjamini-Hochberg multiple test correction.

## Acknowledgements

We thank Drs. Penny Amy, Kurt Regner, Philippos Tsourkas, and Diane Yost for assistance in obtaining the *P. larvae* phages.

## Data Availability

Analysis scripts and processed data were published on Github at https://github.com/jtvanleuven/plarvae_resistance. Genome assemblies and raw sequencing reads for the seven starting phages (SRR31700794-SRR31700800) and the ancestor (starting isolate) of B-3650 (SRR31700744, SRR31700745) were deposited at NCBI under BioProject number PRJNA1130131. Raw sequencing reads (SRR29659251-SRR29659276) for the evolved phage-resistant isolates of B-3650 were deposited under the same BioProject number.

## Supplemental Information

SI Table 1. Cross-resistance testing of all phages on phage-resistant isolates. Sheet “Cross Resistance Streaking” shows qualitative analysis of resistance by streaking bacterial isolates across a line of phages (see methods and SI Figure 1). Sheet “Spot Plating Cross Resistance” shows the results of spot plating 5 uL of phage stock on resistant hosts. The color of the cells indicate level of clearing within the spot. If there were a countable number of plaques within the spot, that number is show. The “Titering” sheet shows the results of tittering the phage stock on different phage-resistant hosts. The titer is shown in pfu/mL. The same phage stock was used in all three experiments.

SI Table 2. Results of efficiency of plating assays. All seven phage stocks were plated on all resistant hosts using the soft-overlay method. Plaques were counted and the phage titers on each host back calculated. These titers are shown in pfu/mL. The first sheet (“Full Spread Sheet”) reports all of the data.

The other spreadsheets are subsets of this larger dataset.

SI Table 3. Output from bacterial growth experiments for 14 different phage resistant isolates. Each isolate was grown in replicates of three. The B-3650 strain as well as the isolate of B-3650 that we picked just prior to challenge (Anc) were both tested. Reported values are OD600 measurements.

SI Table 4. Results showing which phage-resistant isolates has significant growth defects. The effect size on the three measures of growth for significantly different isolates is shown. These values are reported by the R package gcplyr (64).

